# Engineered bacteria launch and control an oncolytic virus

**DOI:** 10.1101/2023.09.28.559873

**Authors:** Zakary S. Singer, Jonathan Pabón, Hsinyen Huang, William Sun, Hongsheng Luo, Kailyn Rhyah Grant, Ijeoma Obi, Courtney Coker, Charles M Rice, Tal Danino

**Affiliations:** Department of Biomedical Engineering, Columbia University, New York, NY 10027, USA; Laboratory of Virology and Infectious Disease, The Rockefeller University, New York, NY 10065, USA; Herbert Irving Comprehensive Cancer Center, Columbia University, New York, NY 10032, USA; Data Science Institute, Columbia University, New York, NY 10027, USA

## Abstract

The ability of bacteria and viruses to selectively replicate in tumors has led to synthetic engineering of new microbial therapies. Here we design a cooperative strategy whereby *S. typhimurium* bacteria transcribe and deliver the Senecavirus A RNA genome inside host cells, launching a potent oncolytic viral infection. “Encapsidated” by bacteria, the viral genome can further bypass circulating antiviral antibodies to reach the tumor and initiate replication and spread within immune mice. Finally, we engineer the virus to require a bacterially delivered protease to achieve virion maturation, demonstrating bacterial control over the virus. This work extends bacterially delivered therapeutics to viral genomes, and shows how a consortium of microbes can achieve a cooperative aim.

## Introduction

The broad range of applications that employ bacteria and viruses for therapy mirrors the diversity of microbes themselves. Distinct bacteria target different tissues, microbiomes, and even intra- versus extra-cellular spaces. Ranging from skin-colonizing *S. epidermidis* to lung-homing *M. pneumoniae*, to the intracellularly growing *L. monocytogenes* and *S. typhimurium*, different bacteria are each equipped with capabilities uniquely exploitable for synthetic engineering ^1–5^. Analogously, the array of viral families under investigation for therapy is similarly broad, with applications exploiting the different genomic structures, natural tropisms, and life cycles of each. Examples include small DNA viruses like AAV for non-immunogenic tissue targeting, minus-strand RNA viruses like rabies virus for retrograde neuronal tracing, and plus-strand RNA viruses like the poliovirus- derivative “PVSRIPO” for oncolytic purposes ^6–10^. Thus, microbes of broadly distinct cellular proclivities have each found utility for a specific application niche.

Bacteria and viruses are generally considered separately in approaches to therapeutic delivery. Yet, during natural co-infection, some viruses and bacteria directly bind to one another, leading to enhanced fitness ^11–13^. Similarly, applications in synthetic biology are beginning to engineer coordination between multiple interacting entities with unique properties that can together produce a consortium achieving a collective objective ^14–21^. In this work, we consider synthetic approaches for viral and bacterial cooperation to overcome key challenges in applying microbially-delivered therapies to combat cancer. While oncolytic viruses have shown efficacy in clinical trials, pre-existing humoral immunity can limit the ability of systemically delivered viral particles to reach their target tumors^22^. Meanwhile, engineered bacteria therapies can deliver genetic payloads to tumor cells, but typically remain trapped locally within the tumor core and are limited in their ability to influence peripheral or distant tumor cells. Indeed, clinical trials with strains such as attenuated *Salmonella* VNP200009 have yet to demonstrate significant clinical efficacy and show dose-related toxicity ^23,24^.

Here, we address these limitations by engineering a platform called CAPPSID (**C**oordinated **A**ctivity of **P**rokaryote and **P**icornavirus for **S**afe **I**ntracellular **D**elivery). This system consists of a synthetic bacteria-virus partnership to deliver an oncolytic picornavirus into tumors via systemic delivery, where the bacteria can cloak the virus from circulating antiviral antibodies. Then, once inside the tumor, the bacteria can launch the virus and spread. Specifically, CAPPSID relies on an engineered *S. typhimurium* to act as a dynamic and synthetic “capsid” to transcribe and deliver viral RNA inside cancer cells, launching a virus that can directly lyse surrounding cells. We then additionally engineer the virus to require a bacterially donated accessory enzyme necessary for viral maturation and subsequent spread, thereby constraining viral replication to one additional infection cycle beyond the originating, bacteria-containing cell (**Fig. 1**). Together, this bacteria-viral cooperation enables tumor inhibition, and yields prolonged engineered viral replication. As the first example of direct engineered cooperativity between bacteria and oncolytic viruses, this work demonstrates bespoke interacting communities of programmable medicines.

**Figure 1.**
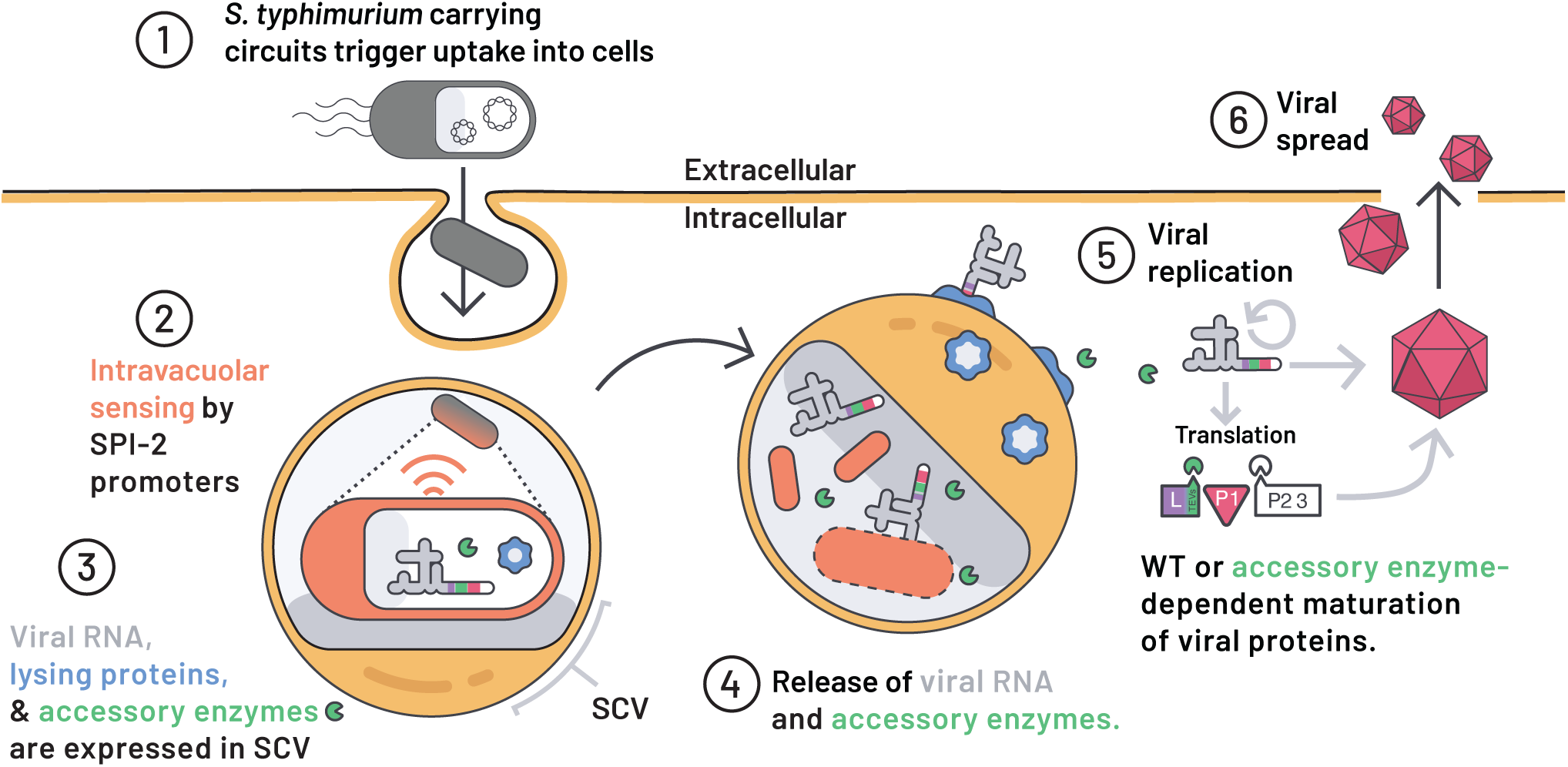
Programmed S. typhimurium autonomously lyse in host cytoplasm to launch viral RNA and an essential orthogonal viral protease. (1) S. typhimurium carrying synthetic circuits enter mammalian cells via natural effectors encoded on Salmonella Pathogenicity Island 1 (SPI-1). **(2)** Internalized S. typhimurium within a Salmonella Containing Vacuole (SCV) sense the intravacuolar space and trigger activation of SPI-2 promoters. **(3)** Engineered SPI-2 promoters are then used to drive the production of viral RNAs (poliovirus replicon, Senecavirus A (SVA), or engineered SVA), lysing proteins hemolysin-E (HlyE) and E from phage φX174, and accessory enzyme. **(4)** Upon successful bacterial and vacuolar lysis, viral RNAs and accessory enzyme are released into the host cytoplasm. **(5)** Wild-type viral RNAs are translated in the cytoplasm and viral replication is initiated. The maturation of viral particles may be engineered to require the accessory enzyme for complete maturation. **(6)** Infectious particles are released into the extracellular space to infect neighboring cells. Since S. typhimurium bacteria act as a viral “capsid”, we have named the platform **C**oordinated **A**ctivity of **P**rokaryote and **P**icornavirus for **S**afe **I**ntracellular **D**elivery (CAPPSID).

## Results

### Engineered *S. typhimurium* autonomously launches viral RNA

To establish CAPPSID as a bacterial platform capable of delivering viral RNAs into cells, we focused on *S. typhimurium*, a naturally facultative intracellular bacterium. *S. typhimurium* achieves invasion into host cells via macropinocytosis and survives within the Salmonella Containing Vacuole (SCV) by expressing a battery of genes encoded on Salmonella Pathogenicity Islands 1 and 2 (SPI-1 and SPI-2), respectively ^25^. By using Salmonella to release viral RNA, the direct translation of its viral gene products bypasses the need for nuclear translocation of plasmid-encoded therapeutics and their subsequent expression before induction of apoptosis or pyroptosis by *Salmonella* ^26,27^. To efficiently deliver long nucleic acids by bacteria, associated challenges include sufficient RNA production, robust RNA integrity, RNA escape into the host cytoplasm, and the need for the host to survive through protein translation.

To control RNA release, we employ natural spatial cues transduced by Salmonella to sense the intracellular phase of their life cycle, using the SPI-2 promoters of the genes sseA and sseJ to trigger activation^28–31^ (**Fig. 2A**). When these promoters are fused to mCherry, we observed rapidly increasing expression of both promoters only after entering mammalian cells, indicating that they are tightly regulated and suitable for intracellular cargo expression, consistent with previous reports (**Fig. 2B,C, S1A**) ^28,32–39^.

**Figure 2.**
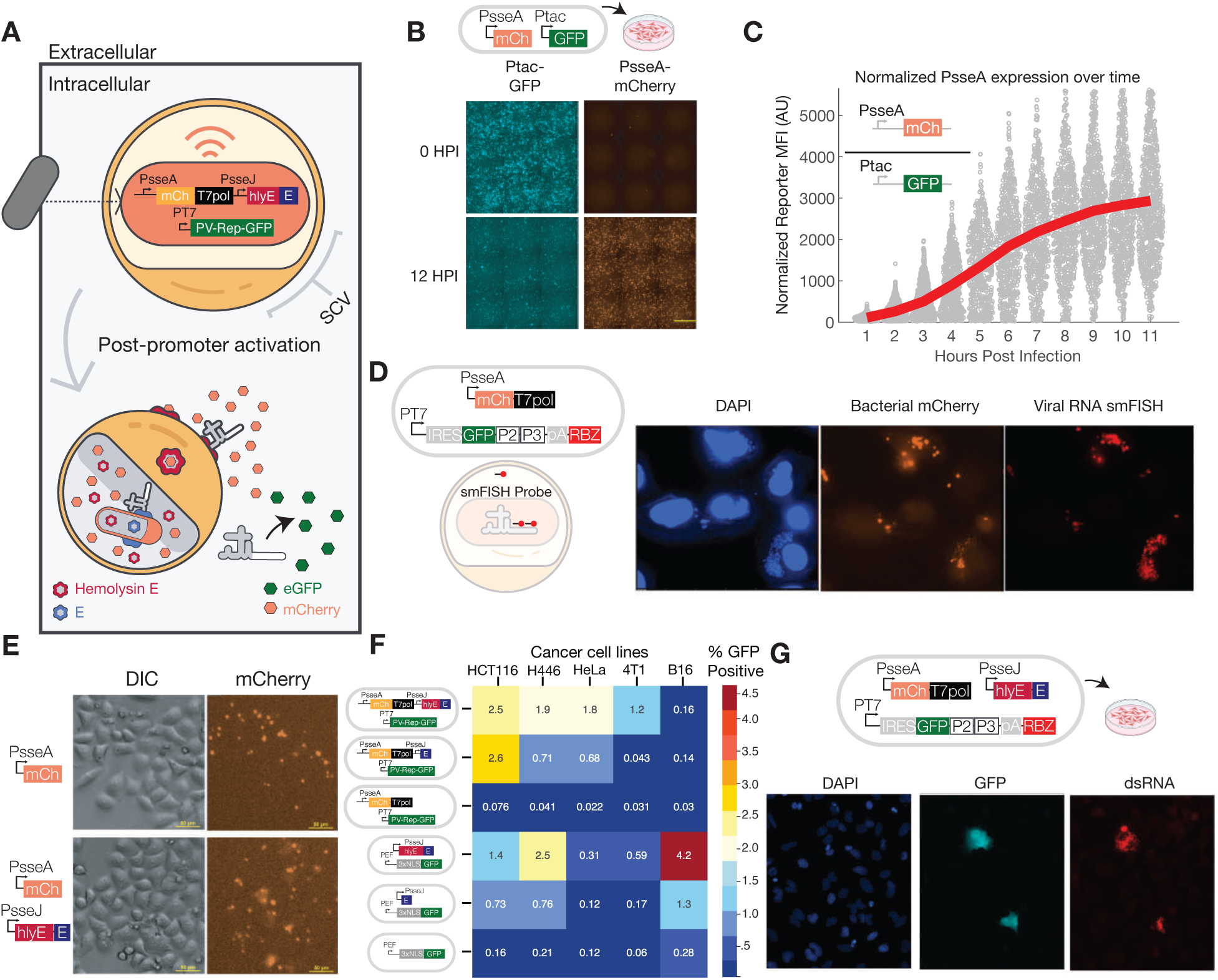
Engineered bacteria deliver self-replicating RNA into the cytoplasm of host cells. **(A)** Intracellular S. typhimurium activate SPI-2 promoters PsseA and PsseJ, to drive mCherry. After internalization of bacterium, PsseA drives mCherry and T7 polymerase, while PsseJ drive the two lysis proteins. As a result, intravacuolar S. typhimurium lyse themselves and the SCV, releasing mCherry and T7-driven poliovirus-replicon RNA into the cytoplasm where replication and translation produce reporter GFP. (**B)** Microscopy images of HeLa cells inoculated with S. typhimurium at multiplicity of infection (MOI) 50 carrying Ptac GFP and PsseA-mCherry plasmids. The top panels show constitutive Ptac-GFP and SCV-induced PsseA-mCherry signals at 0 HPI. The bottom panels show respective signals 12 HPI. Scale bar is 500μm. **(C)** Quantification of PsseA activation is shown as the mean fluorescent intensities (MFI) of mCherry divided by GFP, where each dot represents a single HeLa cell. At each time point the average over all cells is plotted as a red line. The initial value is taken 1 HPI. **(D)** (Left), Top- circuit diagram of proteins produced by PsseA activation and T7-driven poliovirus replicon. Bottom- Schematic of smFISH probes binding specifically to the 3’ end of the viral RNA transcribed by bacteria. (Right) Micrograph showing DAPI staining of both mammalian and bacterial DNA (blue), PsseA-mCherry fluorescence from S. typhimurium inside SCVs (orange) and fluorescent signal from probes specific to viral RNA (red). **(E)** Top panels show representative DIC and mCherry signals of HeLa cells inoculated with S. typhimurium at an MOI 50 carrying a PsseA-mCherry plasmid. Bottom panels are HeLa cells inoculated with S. typhimurium at MOI 50 carrying the mCherry reporter and lysing proteins. The white arrow indicates a cell with an mCherry signal diffusing through the host cytoplasm. Scale bar is 50μm. **(F)** In HCT116, H446, HeLa, 4T1, and B16s, the fraction of GFP-positive cells following inoculation. GFP is expressed from poliovirus replicon (first five rows) or via plasmid delivery (GFP expressed off plasmid with mammalian pEF-promoter) from strains that either lyse with HlyE and E proteins, the E protein alone or do not lyse. **(G)** Bacteria with lysing circuit and virus-encoding plasmid are used to inoculate HeLa cells. DAPI indicates nuclear staining; GFP fluorescence is derived from the poliovirus replicon reporter. Red fluorescence signal is an anti-dsRNA antibody (J2) indicating active replication of viral RNA.

Because prokaryotes do not incorporate 5’m7G caps required for typical mammalian translation ^40^, we chose to deliver viral RNAs that rely instead on cap-independent translation using the internal ribosome entry site (IRES) from Picornaviridae ^41^. To evaluate how such a platform might function across a range of cell lines, we first utilize a poliovirus replicon with GFP in place of its structural proteins as our viral RNA reporter, which has been shown to be replication-competent in numerous cell types ^42^. This replicon, with a fully active viral polymerase, serves to self-amplify the viral RNA, but is unable to spread. Notably, the robust GFP production we observed upon direct RNA transfection was reduced by 50- to 1000-fold when the viral polymerase was mutationally inactivated – supporting the use of self-amplifying RNA for increased cargo expression. (**Fig. S1B**). To couple transcription of this viral RNA production to intracellular sensing, the PsseA promoter of *S. typhimurium* drives expression of the highly processive T7-RNA polymerase, which in turn transcribes the viral RNA from its cDNA genome encoded on a plasmid. When this circuit is transformed into *S. typhimurium* LH1301 (ΔaroA, ΔphoPQ) and used to invade HeLa cells, these bacteria produce the full-length viral RNA, as measured by single-molecule fluorescence *in situ* hybridization (smFISH) using probes against the 3’ end of the 5.5kb poliovirus replicon (**Fig. 2D**).

Once transcribed, the viral genome must escape the bacterium and translocate through the SCV into the cytoplasm of the mammalian host to begin replication. To optimize efficiency of this translocation, we added to our circuit two distinct lytic proteins: Lysis protein E from phage φX174 that disrupts bacterial membranes ^32,43–45^, allowing the viral RNA to exit the lysed bacterium, and Hemolysin E (HlyE) which forms pores in the SCV, allowing the viral RNA to enter the host cytosol ^46^. These genes are expressed under the control of intracellular sensing promoter PsseJ, and complemented by a deletion of the *sifA* gene, which further disrupts

SCV integrity ^32,47^. When *S. typhimurium* carries this circuit into HeLa cells, mCherry appears to diffuse out of the SCV, filling the cytoplasm of the host cell, while in the absence of these lytic proteins, mCherry remains punctate, indicating restricted localization within vacuoles (**Fig. 2E**).

Finally, we coupled the viral transcription and lysis circuits together in *S. typhimurium* to evaluate whether the poliovirus replicon encoding GFP could be delivered into a range of cell types and launch replication. We observed intense GFP signals indicative of successful viral delivery and replication in both mouse and human cell lines including 4T1, B16, HCT116, HeLa, MC38, and H446 cells, with varying efficiencies (**Fig. 2F, S1C**). Furthermore, lysis via E and HlyE enabled dramatically more efficient delivery than in the absence of lytic proteins, or when *S. typhimurium* delivered a plasmid-encoding a mammalian-promoter driven GFP, across most cell lines **(Fig. S1D-E)**. Finally, to confirm that this was not simply passive translation of the incoming genomic viral RNA, we stained cells with an antibody against long double-stranded RNA (dsRNA), a product of active viral replication, and observed positive signals in cells that were also GFP positive (**Fig. 2G**). Together, these data show that CAPPSID is capable of successfully delivering actively replicating viral RNA.

### Delivery of full-length oncolytic virus by CAPPSID clears subcutaneous SCLC tumors

We next tested the ability of this system to deliver a therapeutically relevant full-length oncolytic virus, Senecavirus A (SVA), known to efficiently infect neuroendocrine-like cells, including H446 small cell lung cancer ^48–51^. Because cells infected with *S. typhimurium* frequently die via induction of apoptosis and pyroptosis ^26^, a spreading virus could infect surrounding *S. typhimurium*-free cells, thereby augmenting the overall therapeutic effect (**Fig. 3A**).

**Figure 3.**
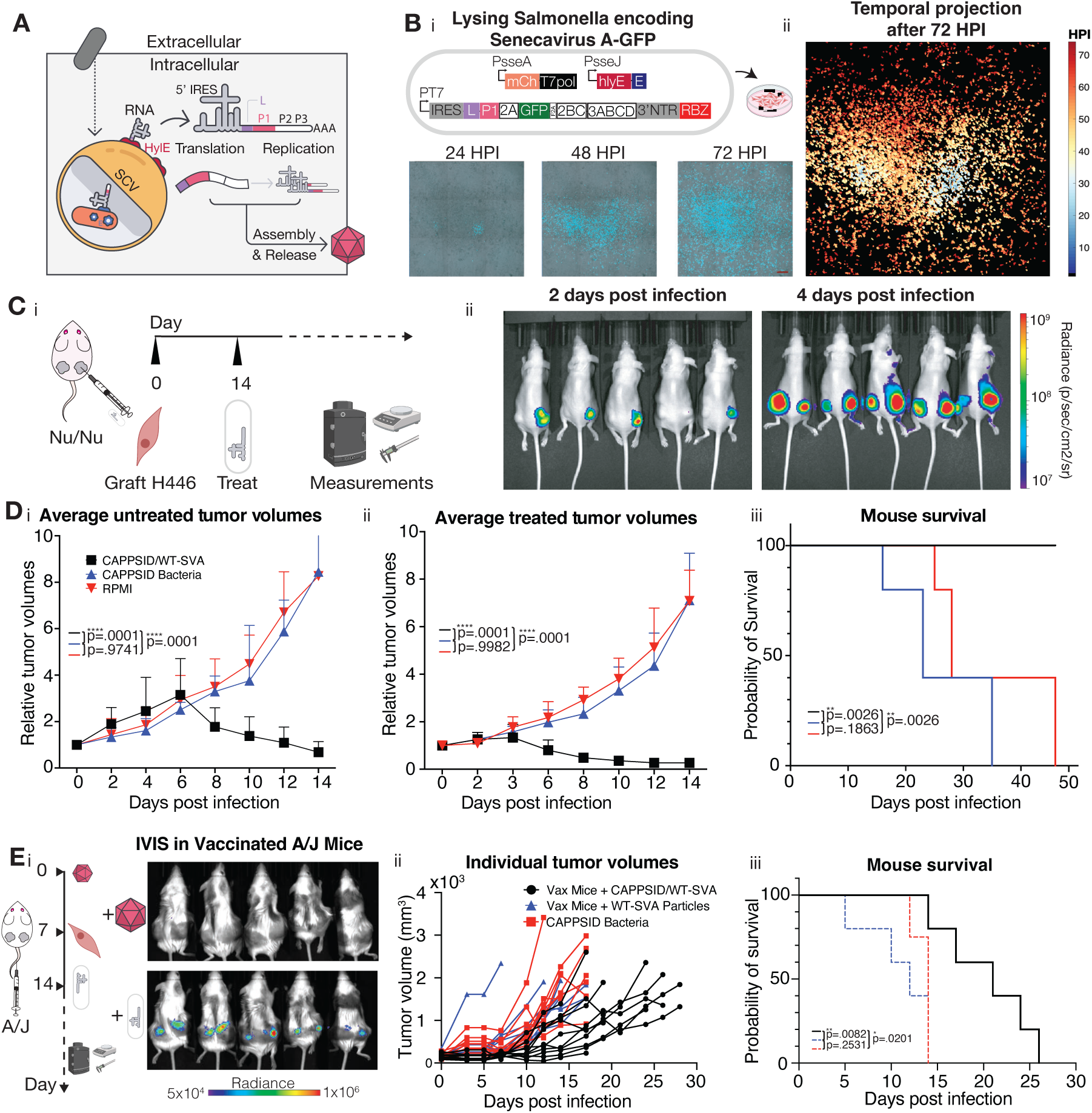
CAPPSID launches full-length oncolytic Senecavirus A. **(A)** Schematic of S. typhimurium SPI-2 driven full length viral RNA and lysing proteins. Upon release of RNA into the host cytoplasm, IRES-mediated translation produces viral proteins necessary for replication of its RNA and virion assembly and packaging. **(B)** H446 cells inoculated with (i, top) MOI 25 S. typhimurium carrying SPI2-driven lysis and reporter plasmid, along with SVA-GFP plasmid. (i, bottom) Time-lapse microscopy at three time-points of spreading SVA-GFP as launched from bacteria. Scale bar is 500“m. (ii) Time course of SVA- GFP infection through the 72-hour acquisition period, projecting time as a color where initial events are represented in light blue hues and later events as yellow and red hues. **(C)** (i) Experimental outline of in vivo experiment where nude mice were engrafted with H446 cells on bilateral flanks, and right flanks were intratumorally injected with 2.5x10^6^ bacteria when tumors reached approximately 150 mm^3^ 14 days later. (ii) IVIS images of nude mice injected with NanoLuc substrate intratumorally two and four days post bacterial injection, demonstrating viral spread. **(D)** (i) Growth kinetics of untreated tumors after administration of lysing S. typhimurium with WT-SVA (black), lysing S. typhimurium only (blue), and RPMI media control (red). Mean relative tumor trajectories plotted with error bars representing SEM of biological replicates (n = 5 tumors per group; ordinary two-way ANOVA with Tukey post-test). (ii) Growth kinetics of treated tumors (n = 5 tumors per group; ordinary two-way ANOVA with Tukey post-test). (iii) Survival curves for mice treated with groups shown in D. Multiple log rank tests with Bonferroni correction, (n = 5 mice per group). **(E)** (i) (left) Experimental timeline where A/J mice were first infected with 2.5x10^6^ SVA plaque forming units (PFU), and then engrafted with N1E-115 cells 7 days later. When tumors reached approximately 150 mm^3^ (after another ∼7 days), mice were injected intravenously with 2.5x10^6^ CAPPSID/nLuc-SVA (Right) IVIS images of mice previously infected with SVA receiving a rechallenge at day 14 with either viral particle rechallenge of 2.5x10^6^ PFU (top) or 2.5x10^6^ CAPPSID/nLuc- SVA (bot) (ii) Growth trajectories of tumors receiving: CAPPSID-SVA (black) in vaccinated mice; lysing control S. typhimurium (red); vaccinated mice rechallenged with an additional 2.5x10^6^ SVA viral particles (blue). (iii) Survival curves for groups in Eii (log-rank test, n=5 mice per group).

For this system to function, bacteria and virus must both be able to infect the target population, with the former not directly inhibiting the latter. Using SVA with a GFP reporter (SVA-GFP) ^52^, we measured whether and how viral spread was affected when introduced one hour after bacterial pre-infection in H446 cells. At 24 hours post infection (HPI) there was no measurable reduction in the spread of virus in the presence or absence of bacteria (**Fig. S2A**).

In *S. typhimurium* containing the lysis circuit, we added SVA-GFP, and inoculated H446 cells to look for successful initial delivery events from bacteria and a subsequent capacity for spread throughout the same culture. Here, GFP-positive cells lacking mCherry would indicate spreading viral infection in cells not originally infected by bacteria, whereas double-positive cells contain both bacteria and virus. The first wave of SVA- infected cells was observed at approximately 8 hours following bacteria inoculation, with viral spreading occurring continuously throughout the subsequent 60 hours. By 72 HPI, effectively all cells were SVA-positive, regardless of initial bacterial infection status (**Fig. 3B, Supplementary Movie 1**). This system demonstrates that even a low fraction of initially infected cells with *S. typhimurium* could deliver and launch the spread of an oncolytic virus across an entire monolayer of cells.

To determine whether CAPPSID carrying the full length SVA virus could also achieve this effect *in vivo*, we used a mouse model engrafted with bilateral hind flank H446 tumors. Right tumors were injected intratumorally (IT) with lysing *S. typhimurium* carrying SVA-NanoLuc (a luminescent reporter) and imaged over time for luminescence. Two days post-infection, right-flank tumors began to show signal. Then, at day four, the signal was additionally observable in left tumors that had not been injected with bacteria, showing productive viral infection in the right tumors and sufficient titer capable of viral translocation to left-flank tumors (**Fig. 3C**). In contrast, control bacteria recombinantly expressing their own luminescent reporter luxCDABE (and no viral RNAs) under the control of PsseA, showed no detectable translocation to left tumors over the same time (**Fig. S2B-C**). Tumor volume measurements over more than 40 days showed complete regression of both left and right tumors in the treatment group within two weeks, whereas all tumors treated with only buffer or lysing bacteria alone continued to grow until reaching maximum allowable sizes (**Fig. 3D, S2D**). The striking regression of tumors conferred 100% survival in mice treated with *S. typhimurium* carrying SVA-NanoLuc, while all control mice treated with RPMI and lysing *Salmonella* alone succumbed to tumor burden. Mice experienced no decline in weight, and negligible bacteria in the liver or spleen, despite appreciable loads present in the tumors, suggesting no adverse response to bacterial injections (**Fig. S2E, F**). Furthermore, histology shows viral spread well beyond the local vicinity in the tumor colonized by bacteria (**Fig. S2G**). Taken together, CAPPSID/nLuc-SVA can launch an oncolytic viral infection clearing H446 tumors *in vivo*.

### Systemic administration of CAPPSID prolongs survival of immunocompetent and SVA-immune mice bearing syngeneic tumors

Thus far, we have shown successful viral launch using IT-injected bacteria in athymic mice unsusceptible to SVA. Next, we sought to extend the investigation of the platform to a fully immunocompetent model, employing the syngeneic mouse neuroblastoma cell line N1E-115 in A/J mice. After validating *in vitro* launch in N1E-115 (**Fig. S2H**), we engrafted mice with double-hind flank N1E-115 tumors, and two weeks later IT injected CAPPSID/SVA. Following the mice over a course of one month, we observed successful attenuation of tumor growth and improved overall survival (**Fig. S2I**).

If CAPPSID can initiate oncolytic infections in tumors when delivered IV, we hypothesized it could also be able to overcome pre-existing immunity against an oncolytic virus. For example, could a bacterially-cloaked viral genome escape circulating antiviral antibodies from a prior vaccination, permitting tumor targeting when systemically delivered? To test this, we first infected mice with WT-SVA particles 7 days prior to tumor engraftment and confirmed the presence of resulting circulating neutralizing antibodies against the virus (**Fig. S2J**). Then, after sufficient tumor growth, we IV treated the mice with CAPPSID/SVA after pre-dosing with anti-innate immune antibodies, and tracked the effects on tumor progression and luminescence over time. We observed that previously vaccinated mice were completely refractory to viral particle rechallenge, resulting in no detectable viral luminescence within the tumor and no survival benefit compared to mock treatment (**Fig. 3E**). In contrast, previously SVA-infected mice that received bacteria in the form of CAPPSID/SVA showed clear luminescence within the tumor as well as improved survival (**Fig. 3E, S2J-K**). Assaying animal health and concurrent biodistribution of these bacteria when delivered IV showed significant enrichment within the tumors, with little to no bacteria detectable in either the spleen or the liver (**Fig. S2L**) and no meaningful decrease in mouse health (weight). Together these data demonstrate the ability of CAPPSID to safely attenuate tumor growth in a fully immunocompetent model when delivered either intratumorally or systemically, even in the presence of neutralizing antibodies.

### CAPPSID enables engineered cooperation between virus and bacteria to control viral spread

While potentially promising, some viruses considered pre-clinically as oncolytics can spread systemically and cause undesirable side-effects^53^. In contrast, the deletion of structural proteins from a virus to yield a so-called replicon or “self-amplifying RNA” has recently gained interest as a therapeutic medium^54^ but still show limited persistence due to an inability to re-infect cells. Methods to improve targeting and safety profiles of oncolytic viruses are achieved either at the cell-surface level, where the receptor-binding domain is altered to recognize a target cell more specifically, or intracellularly, where replication of the virus is modulated positively or negatively by cell-type specific cytoplasmic or nuclear determinants ^55^. Here, we wondered whether we could exploit our CAPPSID platform to control and enhance the virus using the bacteria which are naturally restricted to the immunoprivileged tumor core.

In the life cycle of picornaviruses, all proteins are first translated as a single large open reading frame, termed a polyprotein, that is then cleaved into constituents by virally encoded proteases. These cleaved proteins act as mature replicase proteins and viral structural proteins required for continued spreading infection ^56^. We hypothesized that shifting a cleavage event within the structural proteins to an orthogonal protease expressed by the same bacteria would enable complete viral particles to be produced, leading to a wave of infection beyond the bacterially infected cells, further enhancing the spatial and temporal reach. In those cells subsequently infected by virus and not bacteria, the viral replication could cause cytopathic effects, but spread no further (**Fig. 4A**). Due to its potential for recombinant expression and thorough characterization, we chose Tobacco Etch Virus protease (TEVp) as the orthogonal protease ^57^. Furthermore, TEVp has the flexibility to recognize nearly all residues at the final position of the cognate TEV cleavage site (TEVs) (ENLYFQ^G) where cleavage occurs between the last two amino acids ^57^. This allows for the ability to retain the native N-terminal residue of the downstream protein following successful cleavage.

**Figure 4.**
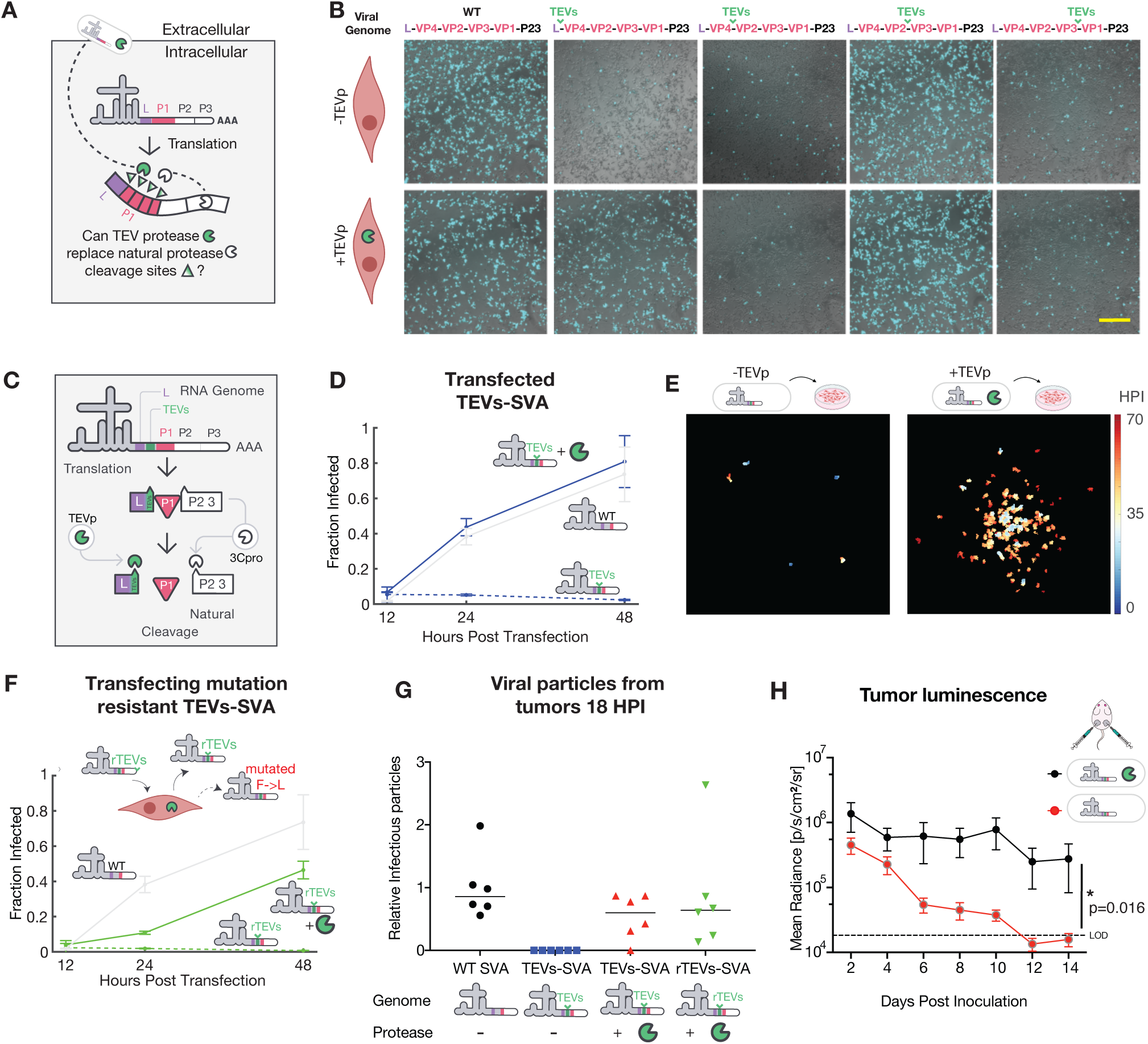
Engineered control over viral protein maturation via CAPPSID-delivered protease. **(A)** Schematic of polyprotein cleavage at natural sites flanking the structural viral proteins as possible sites to reprogram for exogenous TEV protease recognition (green-to-white graded arrows). **(B)** H446 cells stably expressing TEV protease (bottom row) and wild-type H446 cells (top row) were subsequently transfected with 500 ng of WT-SVA and SVA RNA engineered to contain TEV cleavage sites in place of native recognition sequences between structural proteins. Images taken 24 hours post-transfection at 10x. Scale bar is 500μm. **(C)** Illustration of TEV-mediated cleavage between L-protein and VP4. TEV site is cleaved by S. typhimurium-provided TEV protease, leaving a 6 amino acid C-terminal addition (ENLYFQ) on the L protein, while all other cleavage sites are naturally cleaved by the SVA-protease 3Cpro. **(D)** Fraction of GFP-positive cells after transfection of WT SVA-GFP (grey), TEVs-SVA-GFP into H446 cells not expressing protease (dashed blue), and TEVs-SVA-GFP into H446 cells expressing TEVp (solid blue) at 12, 24, and 48 hours. **(E)** H446 cells are inoculated with S. typhimurium carrying TEVs-SVA- GFP with (right) and without (left) TEVp at MOI 50, followed by time-lapse imaging. Temporal information is shown as a projection using color, with initial events represented in light blue hues and later events as yellow and red hues over 72 hours. **(F)** Same as in 4E but with mutationally resistant TEV site (rTEVs, ENLY**C**Q^G). **(G)** Normalized viral titer from bacterially delivered SVA-variants with or without protease. Hind flank H446 tumors IT injected with lysing S. typhimurium carrying WT-SVA-NanoLuc without TEV protease, TEVs-SVA- NanoLuc with and without TEV protease, and rTEVs-SVA-NanoLuc with TEV protease were harvested 18 HPI. Then, naive H446 cells were inoculated with the clarified freeze-thawed tumor homogenates to isolate viral particles. Each point represents one tumor. Data is normalized to the mean of the group receiving bacterially delivered WT-SVA. Two-way ANOVA evaluation determined no significant difference (p=0.46) in means in the groups receiving WT-SVA-NanoLuc, TEVs-SVA-NanoLuc with TEV protease, and rTEVs-SVA-NanoLuc with TEV protease. **(H)** Luminescent signals from nude mice with bilateral hind flank tumors injected with S. typhimurium delivering rTEVs- SVA-NanoLuc with (black) and without (red) TEV protease. Each point on the graph illustrates the mean and standard deviation of the luminescent signal of n=10 tumors at each time point with significant difference between groups (Wilcoxon signed-rank test, two-tailed).

Investigating which natural cleavage sites might be amenable to TEVs substitution, we replaced all four cleavage sites flanking the structural proteins, abrogating the natural cleavage sequence in the process. We engineered each of these four potential variants and transfected them either into wild-type H446 cells, or H446s constitutively expressing TEVp, and looked for conditional spreading (**Fig. 4B,C**). When the TEVs was placed between the nonstructural Leader protein and the first structural protein, VP4, the spreading of this variant became entirely dependent on TEVp and was capable of infecting surrounding cells at a rate equivalent to wild-type virus (**Fig. 4D**). Replacing the other natural cleavage sites with TEVs, however, resulted in TEV- independent spread or abrogation of spread entirely.

To couple the spread of this variant to co-infecting bacteria, we engineered lysing *S. typhimurium* to also express TEVp under the control of a second PsseA promoter. Additionally, we incorporated a series of mutations in TEVp previously shown to improve the solubility of the protease ^58,59^ (**Fig. S3A**). When the bacteria delivered a TEVp-dependent virus without bacterially produced TEVp, the virus launches but then fails to spread, as expected (**Fig. 4E, left**). However, when the *S. typhimurium* simultaneously delivered both the TEVp- dependent virus as well as TEVp, localized foci of spreading infection appeared (**Fig. 4E, right**).

We next proceeded to evaluate this platform *in vivo,* first aiming to characterize the stability of the engineered genome, owing to the high error rate of RNA-dependent RNA polymerases (RdRps) and potential to mutate away from TEVp-dependence ^60^. Tumors were injected IT with *S. typhimurium* delivering wild-type SVA- NanoLuc and compared to TEV-dependent virus (TEVs-SVA-NanoLuc) with co-delivered protease. Over the course of one week, the luminescence of the group receiving bacterially delivered wild-type virus continued to increase rapidly. In contrast, the signal from tumors injected with bacterially delivered TEV-dependent SVA along with protease remained lower than wild-type through day eight, but then began to increase (**Fig. S3B, top**). Sequencing viral RNA extracted from tumors that received the TEVs-SVA-NanoLuc revealed that three out of five had a single-nucleotide polymorphism (SNP) in the TEVs in >95% of total reads, yielding two different ways of producing an identical phenylalanine-to-leucine (F->L) substitution (**Fig. S3B, bottom**). When this mutation was cloned into the viral genome and transfected directly into H446 with or without TEVp, we observed that this mutation was indeed sufficient to achieve TEVp-independent spreading (**Fig. S3C**). Examination of the natural cleavage site at this position revealed that having a leucine at this site recapitulates the amino acid normally present immediately upstream of the native scissile Q^G site.

To prevent this “escape” mutation from occurring, an optimal TEVs sequence would be one where the codon for phenylalanine requires more than one SNP to revert into a leucine. While no codon like this for phenylalanine exists, previous interrogation of TEVs revealed that a cysteine substitution at the phenylalanine site maintained TEVp-mediated cleavage, while also being two SNPs away from mutating to a leucine ^61^. Indeed, an SVA variant with the modified TEVs sequence of ENLY**C**Q^G only spread conditionally in the presence of TEVp, though at slightly reduced efficiency compared to the WT TEVs (**Fig. 4F, S3C**). Thus, we were able to construct a mutationally resistant variant of TEVp-dependent SVA (denoted rTEVs-SVA).

Mice carrying double hind flank H446 tumors were injected IT with CAPPSID rTEVs-SVA with and without TEVp expression, or WT-SVA alone. Then, 24 hours following injection of bacteria, tumors were harvested, homogenized, and assayed for luminescence *ex vivo* as a readout for replication originating from bacterial launch, as well as for viral titer measurements as an indication of successful packaging of the virus. The initial luminescence as measured *ex vivo* was statistically indistinguishable between groups, showing equivalent initial delivery from bacteria of WT virus compared to TEVp-dependent virus (**Fig. S3D**). Similarly, the number of viral particles produced by cells infected with TEVp-dependent virus was also statistically the same as WT virus delivery at launch (**Fig. 4G**). In contrast, no infectious particles were recovered when the TEVp-dependent virus was delivered in the absence of TEVp, as expected (**Fig. 4G**). Furthermore, when naive cells *in vitro* were infected with tumor-harvested WT virus, spreading was observed, while tumors containing TEVp-dependent virus showed initial replication, and no further spread (**Fig S3E**). Together, these data suggest that the initial launch and production of infectious viral particles are equally efficient between both WT and TEV-dependent viruses, and that engineered virus launched from bacteria was indeed TEVp-dependent. Finally, when a cohort of mice injected with lysing *S. typhimurium* delivering TEVp-dependent virus with and without protease was measured longitudinally, we observed that luminescence from this mutation-resistant variant continued to remain present for two weeks following a single injection, while virus delivered without protease showed a complete loss of signal over the same period. Over this time course, no increasing luminescent signals were observed, suggesting that reversion to TEVp-independence did not occur (**Fig. 4H, S3F**).

## Discussion

This work explored synthetic strategies to design distinct levels of cooperation between two clinically relevant microbes - *S. typhimurium* and Senecavirus A. By utilizing bacteria as a dynamic and engineerable “synthetic capsid”, we delivered replicons and full-length viral RNAs into the host cytoplasm. Using CAPPSID, we entirely clear subcutaneous SCLC tumors and overcome systemic neutralizing antibodies when injected IV in fully immunocompetent mice. Additionally, the bacterial launch of replicons into a range of mouse and human cell lines demonstrates the ability to deliver non-spreading self-amplifying viral RNAs beyond their natural tropism. Noting that the bacteria can simultaneously deliver proteins and nucleic acids, we devised further interactions by engineering a virus whose protein maturation depends on a bacterially provided protease when substituting a synthetic cleavage site. Together, we developed a multi-layered engineering approach for coordinating a two-microbe system for oncolytic applications. The platform was further able to show how coordination can dramatically improve the persistence of the virus compared to a replicon when supplemented with an engineered requirement for an accessory enzyme for viral spreading. While SVA does not cause mouse toxicity, the development of new strategies to achieve targeted replication and overcoming systemic neutralization have remained central challenges in the advancement of novel virotherapies. Through these efforts, our CAPPSID system demonstrates the ability of viruses to extend the tumoricidal effect of bacteria.

Delivery of nucleic acids by bacteria, or bactofection, has been previously applied, for example by *Agrobacterium* for CRISPR/CAS9 gene-editing in wheat ^62^, and *L. monocytogenes*, *E. coli,* and *S. typhimurium* for siRNAs, short ORF-containing RNAs, and plasmids ^27,36,63–71^. Leveraging the flexible tools available for genetic engineering in *S. typhimurium*, we were able to deliver large viral RNAs across a broad range of cell types compared to those previously reported via engineered *S. typhimurium*^66^. Building on previous work for intracellular delivery, the active replication of viral RNA causes a cytopathic effect in its initial host cell, while also enabling spread to surrounding cells uninfected by bacteria, thereby enhancing therapeutic range. While SVA does not natively infect nude mice, we additionally show efficacy in a systemically delivered, fully immunocompetent model susceptible to SVA ^48^. This consortium of cooperative microbes likely elicits its effects through a range of mechanisms, including direct cytopathic effect, innate immune activation via microbial pathogen- and damage- associated molecular patterns, as well as neoantigen cross-presentation in the context of an intact adaptive immune system.

Our efforts to insert an orthogonal cleavage site into the virus highlight the importance of addressing the mutability of RNA viruses. RNA-dependent RNA polymerases incorporate an incorrect base at a rate of roughly 1 in 10,000 ^72–75^. Here, we attempted to mitigate mutational escape by first identifying the most common escape mechanism *in vivo*, and limiting the likelihood of such reversion by requiring two independent mutations to simultaneously occur, thereby geometrically reducing the escape probability. However, alternate types of mutations, such as wholesale deletions of our orthogonal sequence may also occur, though were not observed here. Insertion of additional TEV sites, or even additional protease/cleavage site pairs, could further increase the robustness of this system and enable construction of logic-gated viral replication and spread.

By developing a bacterially delivered platform for viral RNA, we show successful launch of a viral infection capable of eradicating tumors, the ability to cloak and deliver viral genomes into tumors in mice with humoral immunity, and that viral spreading controlled in *trans* by a bacterially donated protease can enhance persistence compared to a replicon alone. Together this engineered microbial consortia produces a potent therapy that overcomes the limitations of singular approaches.

## Methods

### Bacterial cell growth

All bacteria were grown in LB Lennox broth. For LH1301, cultures were supplemented with ampicillin (100 μg/mL) and spectinomycin (200 μg/mL) for viral-encoding plasmids and lysis circuit plasmids, respectively. Plasmids were cloned into NEB10β and maintained at 100 μg/mL ampicillin or 100 μg/mL spectinomycin. To prepare strains of LH1301 containing these plasmids, the cells were electroporated, recovered, and plated on LB agar. All liquid cultures were grown overnight in a 30℃ incubator with shaking. For all *in vitro* and *in vivo* experiments, cultures were grown overnight at 30°C to stationary phase, and then diluted 1:100 the next morning and grown for ∼3h at 37°C until reaching an OD600 of 0.5.

### Plasmid Construction

Plasmids were constructed using HiFi Assembly (NEB NEBuilder HiFi DNA Assembly Master Mix) and transformed into NEB10β. SVA-containing vectors were cloned into the same p15A/amp backbone flanked by a 5’ T7 promoter, and 3’ polyA tail, HDV ribozyme, and T7-terminator. All lysis circuits were constructed by synthesizing gBlocks from IDT, and cloned into plasmids with SC101 origins of replication. To validate constructs, whole plasmid sequencing was performed by Plasmidsaurus.

### Bacterial genome engineering

The *sifA* gene was deleted using the λ-Red recombination system^74^. Linear DNA containing CmR flanked by FRT sites were PCR amplified using pkD3 plasmid and electroporated into LH1301 carrying pKD46 plasmid. Chromosomal deletions were verified by PCR and Sanger sequencing.

### Mammalian cell maintenance

HeLa (CRL-1958), 4T1 (CRL-2539), B16 (CRL-6475), HCT116 (CRL-247), N1E-115 (CRL-2263) and H446 (HTB-171) were acquired from ATCC, and MC-38 from Kerafast (ENH204-FP). All cell lines were cultured in RPMI supplemented with 10% Fetal Bovine Serum (FBS, Gibco) and 1X MEM Non-Essential Amino Acids (NEAA) in a 37°C tissue culture incubator with 5% CO2. To stably and constitutively express TEVp in H446, cells were selected and regularly grown with 0.25ug/ml puromycin after transfection with PB-PEF-Puro-F2A-TEVp and piggyBac super-transposase.

### S. typhimurium in vitro invasion assay

Cultures of *S. typhimurium* were grown to stationary phase overnight and diluted 1:100 the following morning into LB Miller supplemented with antibiotics. Cultures were grown at 37°C to OD600 of 0.4-0.6, when they were spun down and resuspended in 1ml RPMI+1%FBS. Bacteria were added at an MOI 50 into 24 well plates of mammalian cells split 24h prior, and spun down in the plate at 200xG for 5 min. Plates were then incubated at 37°C, 5% CO2 for 30 min. Then, the wells were thoroughly washed with RPMI to remove bacteria, incubated for another 30 min at 100ug/ml Gentamicin in RPMI+10%FBS to kill residual extracellular bacteria, and then replaced with 25ug/ml Gentamicin in RPMI+10%FBS for the duration of the experiment. To stain nuclei, NucBlue (Invitrogen R37605) was added at 10-20ul per ml of media.

### Viral transfection and production

To produce virus, H446 cells were transfected using Lipofectamine MessengerMax at 1ul reagent per 500ng RNA per well in a 24 well plate, and scaled up as needed. 48 hours post transfection, cells were harvested, freeze-thawed three times to disrupt membranes and release viral particles, and clarified by centrifugation at 16,000xg for 10 minutes. Virus was titered on H446 by using serial dilutions and identifying the concentration of virus infecting roughly 50% of cells in a well, plated at 100,000/cm^2^ 24h prior, by imaging luminescence or GFP reporter signal, defined as MOI 1.

### Animal tumor models

All animal experiments were approved by the Institutional Animal Care and Use Committee (Columbia University, protocol AC-AABQ5551). Female nude mice aged 6-8 weeks from Charles River were grafted with bilateral subcutaneous hind flank tumors of 5x10^6^ H446 cells in 50% reduced growth factor Matrigel (Corning). Tumors were grown until reaching ∼150mm^3^ over ∼2.5 weeks. Alternatively, female A/J mice aged 6-8 weeks from Jackson Labs were grafted with subcutaneous hind flank tumors of 5x10^5^ N1E-115 cells in 50% reduced growth factor Matrigel (Corning), and grown for 10 days until reaching ∼150mm^3^. Then, 2.5x10^6^ bacteria in 25ul RPMI (without phenol red) were injected intratumorally, or by IV in 100ul. For IV delivery of bacteria in A/J mice, subjects also received 250ug each of IL1R-, TNF⍺-, and IFNAR1-antagonizing antibodies 2h prior to bacterial injection (BioXCell BE0256, BE0058, BE0241). For IVIS imaging, mice were sub-tumorally injected with 25ul Nano-Glo Fluorofurimazine In Vivo Substrate (Promega N4100) at 8.8 nM. Tumor volume was quantified using digital calipers to measure the length and width of each tumor (V=0.5*L*W^2^). The protocol requires animals to be euthanized when tumors reach 2cm in diameter or under veterinary staff recommendation. Survival curves were generated based on tumor burden as all mice survived engraftment and were not directed for euthanasia before reaching tumor burden limit. Mice were randomized into various groups in a blinded manner.

### Biodistribution

After time points following bacterial injections indicated in figure legends, mice were euthanized to collect the tumors, spleen, and liver. Tissues were weighed and homogenized using a gentleMACS tissue dissociator (Miltenyi Biotec; C-tubes). These homogenates were then 10-fold serially diluted and plated on LB agar with chloramphenicol and grown overnight at 37°C. Colonies were counted and computed as cfu per gram of tissue.

### *Ex vivo* tumor luminescence

Tumors were extracted, weighed and homogenized using the gentleMACS tissue dissociator (Miltenyi Biotec; C-tubes) in 5 ml RPMI+1%FBS. To measure luminescence, samples were serially diluted 10-fold over 4 orders of magnitude, in replicate, and assayed by plate reader (Tecan Infinite 200 Pro using the i-control software version 3.9.1.0) after adding Nano-Glo In vivo Substrate (Promega CS320501).

### *Ex vivo* tumor-associated virus sequencing

Tumors were extracted, weighed and homogenized using a handheld tissue grinder in 2 ml RPMI supplemented with Ambion RNALater (ThermoFisher #AM7020). Total RNA was extracted using the RNEasy kit from Qiagen (#74104). Then, cDNA was prepared using Superscript III First Strand Synthesis Supermix (ThermoFisher #18080400) kit with a poly-dT primer. This output was as PCR template with primers against the SVA genome flanking the TEVs site. The linear ∼500bp fragment was read using Nanopore sequencing provided by Plasmidsaurus, with raw reads aligned using Geneious Prime.

### Ex vivo tumor viral titering and antibody neutralization

Tumors were extracted, weighed and homogenized using the gentleMACS tissue dissociator (Miltenyi Biotec; C-tubes) in 1 ml RPMI+1%FBS with anti-anti (Gibco). Homogenate was freeze-thawed three times and clarified by centrifugation at 16,000xg for 10 minutes. Three 10-fold serial dilutions of this homogenized preparation were inoculated on naive H446 cells for 1hr, and then replaced with fresh RPMI+10%FBS. 12h later, cells were dissociated by pipetting and imaged to count the number of luminescent cells after adding Nano-Glo In vivo Substrate (Promega N4100).

For neutralization assays, mice were cheek bled two weeks after receiving either 2.5x10^6^ viral particles IV or sham. Serum was then isolated by centrifugation, and diluted 1:100 with MOI 1 SVA-GFP in 200ul of 1% FBS in RPMI with Pen/Strep. This inoculum was overlaid on H446s plated at 75,000/cm^2^ 24h prior. After 30min, the inoculum was removed, and then imaged to observe the number of reporter-positive cells.

### Histology

For histology experiments, tumors were excised and fixed in 4% paraformaldehyde, then switched to 70% ethanol, bisected, paraffin-embedded, sectioned at 4um, and sent for histology services at Histowiz. Sections were stained using Anti-Lipopolysaccharide (WN1 222-5, Absolute Anitbody, Ab00141-23.0) or TUNEL.

### Statistical analysis

Statistical tests and corresponding plots were performed and corresponding plots created using either GraphPad Prism 10 or Matlab 2023A. The details of the statistical tests are indicated in the respective figure legends. Mice were randomized into different groups before the experiments.

### ImmunoFISH sample preparation and staining

The ImmunoFISH sample preparation and staining protocol was performed as in Singer et al, 2021 ^76^. Cells were fixed in 4% formaldehyde for 5 minutes min at room temperature, washed twice in PBS, and placed in 70% ethanol overnight at −20°C. For staining dsRNA, the next morning, cells were washed twice with PBS; permeabilized in 0.1% TritonX for 5 minutes min; washed twice in PBS; blocked in 10% Normal Goat Serum (Thermo 50062Z); washed twice with PBS; incubated with J2 primary-antibody (Scicons 10010200) at 0.5ug/ml in 10% Normal Goat Serum for 2 hours; washed twice with PBS; incubated with goat anti-Mouse Alexa Fluor 647 secondary antibody (Invitrogen A-21235) at (room temperature) RT for 30 minutes; washed twice in PBS and incubated with 10% Normal Goat Serum for an additional 10 minutes at RT, and then imaged in PBS.

For smFISH staining, probes were designed to bind the 3’ end of the poliovirus replicon. After fixing and overnight incubation in 70% ethanol, cells were washed twice in PBS; equilibrated in FISH Wash Buffer containing 2X SSC (Invitrogen 15557044) and 20% Formamide (Ambion AM9342) for 5min at RT; and hybridized with Stellaris FISH probes labeled with Quasar 670 at 125nM (Biosearch Technologies, Supplementary Table 1) overnight at 30°C in Hybridization Buffer (containing 20% Formamide (Ambion AM9342), 2X SSC, 0.1g/ml Dextran Sulfate (Fisher Sci BP1585–Dextran Sulfate), 1mg/ml *E. col*i tRNA (Roche 10109541001), 2mM Vanadyl ribonucleoside complex (NEB S1402S), and 0.1% Tween 20 (VWR 97062-332) in nuclease free water). The next morning, the hybridization buffer was removed and cells were washed twice in FISH Wash Buffer; incubated in FISH Wash Buffer without probe for 30min at 30°C; washed three times with 2X SSC; counterstained with DAPI; and finally imaged in 2X SSC.

### Microscopy

Cells were imaged on a Nikon Ti2 with PFS4, a Nikon Motorized Encoded Stage, Lumencor SpectraX Light Engine, custom Semrock filters, and a Prime 95B sCMOS camera. Automated acquisition for snapshots and time-lapse was programmed in NIS Elements. The scope was equipped with an OKO stage top incubator with temperature-, humidity-, and CO^2^-control, enabling long-term imaging. For imaging smFISH a 60x, objective was used. Otherwise, imaging was performed using a 10X or 20X ELWD objective.

### Imaging Analysis

Images were recorded and processed using NIS-Elements software, Fiji, MATLAB, and Python. Time-lapse and smFISH image analysis were performed as in Singer et al, 2021^75^.

### Data availability

The main data supporting the results in this study are available within the paper and its Supplementary Information.

### Reporting Summary

Further information on research design is available in the Nature Research Reporting Summary linked to this article.

## Acknowledgements

We are grateful for input and infectious cDNA clone of SVA from JT Poirier, NYU Langone. T.D. discloses support for the research described in this study from the Department of Defence [BC160541], and National Institutes of Health [R01EB029750]. Z.S.S. discloses support for the research described in this study from National Institutes of Health [F32CA225145].

## Author contributions

Z.S.S. and T.D. conceived of the research. Z.S.S., J.P., H.H., W.S., K.R.G., H.L., I.O., and C.C. performed experiments. Z.S.S., J.P., and T.D. wrote the initial manuscript with input from all authors. Z.S.S. and W.S. performed all revision experiments and manuscript edits. T.D. and C.M.R. supervised the research.

## Supplementary Figures

**Figure S1.**
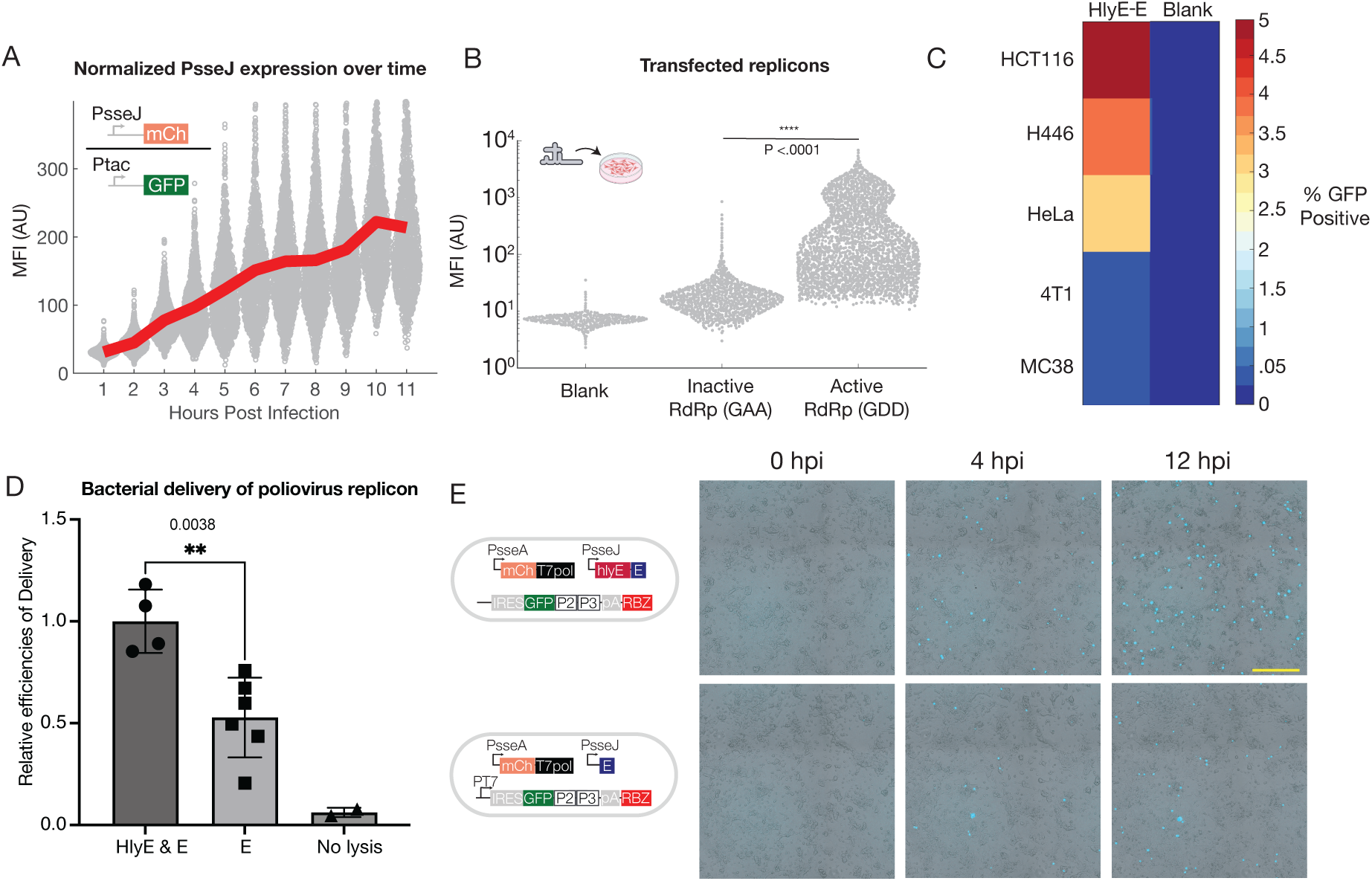
Related to. Figure 2**, Engineered bacteria deliver self-replicating RNA into the cytoplasm of host cells. (A)** Quantification of PsseJ activation is shown as the mean fluorescent intensities (MFI) of mCherry divided by GFP, where each dot represents a single HeLa cell. At each time point the average over all cells is plotted as a red line. The initial value is taken 1 HPI. **(B)** HeLa cells seeded at 20,000/cm^2^ were transfected with replicons containing wild-type polymerase, mutated inactive polymerase (GAA), or mock control. HeLa cells were assayed 18 hours post-transfection and the MFI of each cell is indicated by a single gray dot. From left to right: Blank (AU Mean±SD) 7±1, (Total cell count) n=22,401; GAA polymerase variant 27±28, n=14,666; GDD polymerase variant 1184±1044, n=14960. P<0.0001, KS test. (**C)** Fraction of cells expressing GFP in HCT116, H446, HeLa, 4T1, and MC38 cells indicative of poliovirus replicon replication as delivered by S. typhimurium expressing HlyE and E. **(D)** Comparison of delivery efficiencies with and without HlyE. H446 Cells seeded at 125,000/cm^2 were inoculated the next day with S. typhimurium carrying poliovirus replicon with and without the lysing circuit, HlyE+E, or only E. Efficiency per well is divided by the average delivery efficiency in the HlyE+E group. Statistical test is an unpaired T-test (p=0.0038) **(E)** Time-lapse microscopy images of HeLa cells seeded at 25,000/cm^2^ and inoculated with S. typhimurium carrying either lysing circuit HlyE+E (top) or E only (bottom), both with poliovirus replicon. Scale bar is 500!m.

**Figure S2.**
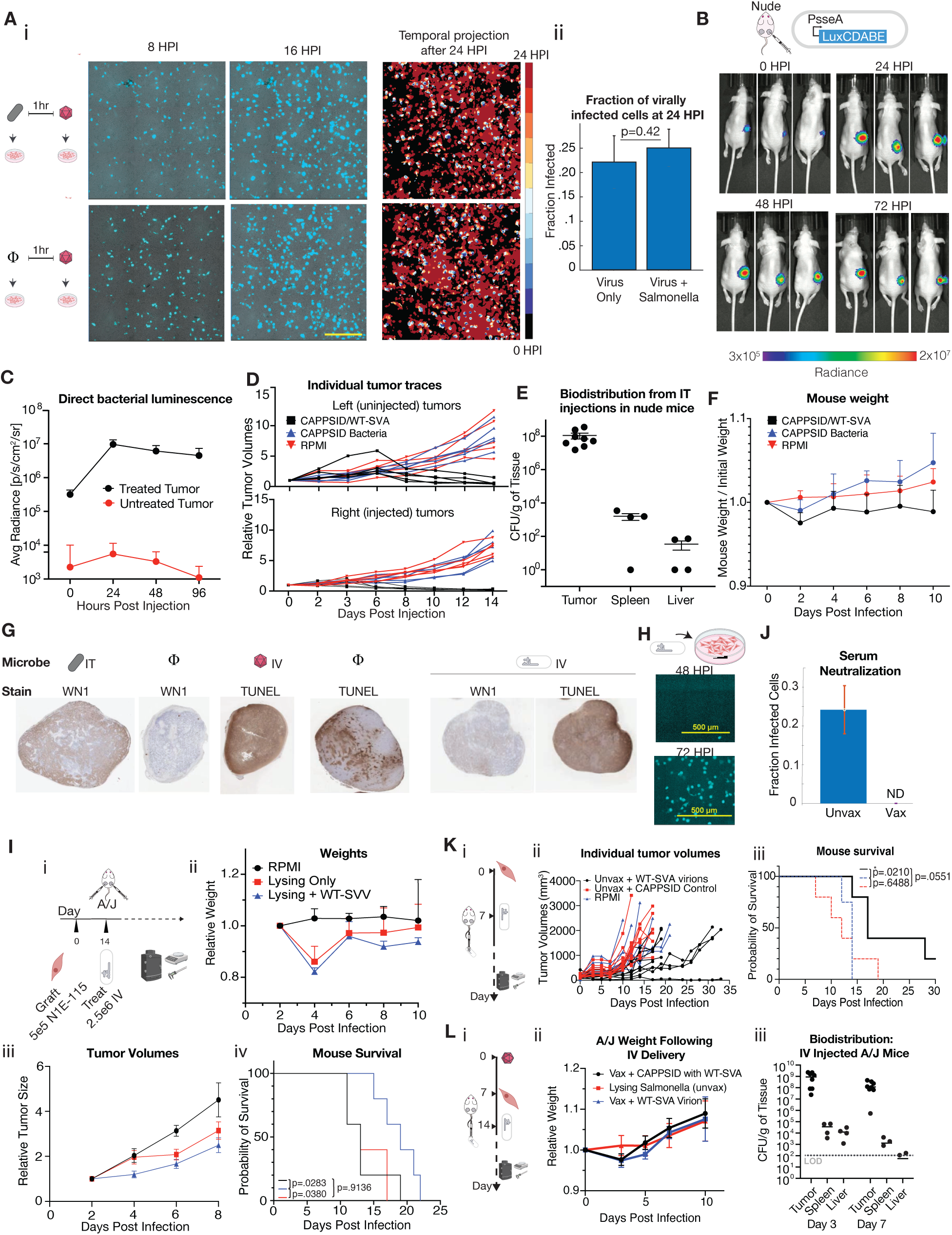
Related to. Figure 3**, CAPPSID launches full-length oncolytic Senecavirus A (A)** (i) Time lapse imaging showing spread of virus in the presence or absence of bacteria. H446 cells seeded at 125,000/cm^2^ were incubated with SVA-GFP at MOI 1 for 20 min (top) or preincubated with S. typhimurium “sifA at MOI 25 for 1h prior to SVA-GFP incubation (bottom). SVA-GFP-infected cells over 24 hours were temporally projected onto a single image for each condition; initial infected cells are represented in light blue with subsequent events depicted as darker shades of red. Movie was recorded over a 24-hour period. Scale bar is 500μm. (ii) Plot depicting fraction of cells infected with SVV at 24hpi when infected with virus alone (left, n=2479), and bacteria infected at MOI 5 1h prior to viral infection (right, n=4108). T-test derived p=0.42 indicates no significant difference in the fraction of replicon-positive cells under both conditions. Scale bar is 500μm. **(B)** In their right-flank tumors, Nude mice received intratumorally 3x10^6^ S. typhimurium expressing PsseA-driven LuxCDABE, and were measured by IVIS over time to identify bacterial translocation. **(C)** From the tumors shown in B, the radiance of left (red) and right (black) tumors were recorded over the first 4 days and reported as average radiance [p/s/cm^2/sr]. **(D)** (i) Nude single mouse trajectories of tumor volume over time. Black lines are mice receiving PsseJ-driven intracellular lysing S. typhimurium with WT-SVA-Nanoluc. Blue lines are mice receiving PsseJ-driven intracellular lysing S. typhimurium only. Red lines are mice receiving RPMI. **(E)** Biodistribution results are represented as CFUs/gram of tissue from the spleens, livers, and tumors of mice 7 days after inoculation with CAPPSID. **(F)** Mouse weights normalized to initial weight over the first 10 days post-inoculation. **(G)** Histology of subtumorally H446**-**engrafted nude mice. From left to right, tumors were treated with 2.5e6 empty CAPPSID bacteria stained for LPS using the WN1 antibody; mock treated and stained with WN1; IV injected with 2.5e6 infectious SVA viral particles and stained for overall cell death by TUNEL; mock treated and stained with TUNEL; and finally, IV injected with 2.5e6 CAPPSID/SVA and then serial sections stained for bacteria with WN1 and TUNEL for apoptotic cell death. **(H)** Schematic of S. typhimurium carrying lysing circuit and WT-SVA-NanoLuc inoculated on N1E-115 cells. Luminescence was captured by microscopy at 48 and 72 hours post infection after addition of luminescence substrate to identify viral launch and spread. **(I)** (i) Illustration summarizing in vivo hind flank engraftment of 5x10^5^ N1E-115 cells on A/J mice and intratumoral injections with lysing S. typhimurium 14 days post-tumor engraftment. (ii) Weight trajectories for mice treated with RPMI (black), lysing Salmonella (red), and lysing Salmonella carrying WT-SVA-NanoLuc (blue). (iii) Tumor volumes over the first 10 days. (iv) Survival curves for treated mice; p=0.028 for S. typhimurium + WT-SVA relative to RPMI and p=0.038 for lysing Salmonella relative to S. typhimurium + WT- SVA. **(J)** Mice were infected with 2.5x10^6^ PFU SVA and two weeks later cheek-bled to collect serum. Serum from naïve mice (“unvax”) and from vaccinated (“vax”) mice was diluted 100-fold and incubated with SVA and N1E-115 cells at an MOI of 1. Bar plot shows the fraction of cells infected with reporter virus 12h later. (N=4 technical replicates) **(K)** (i) Experimental timeline where A/J mice were engrafted with N1E-115 cells. When tumors reached approximately 150 mm^3^ (after another ∼7 days), mice were injected intravenously with 2.5x10^6^ WT-SVA-NanoLuc. (ii) Growth trajectories of tumors receiving: WT-SVA-NanoLuc (black), lysing Salmonella (red), or RPMI media control (blue) (two-way ANOVA with Šídák’s post-test, n=5 mice per group). (iii) Survival curves for groups in Ki (log-rank test, n=5 mice per group). **(L)** (i) Experimental timeline where A/J mice were first infected with 2.5x10^6^ SVA plaque forming units (PFU), and then engrafted with N1E-115 cells 7 days later. When tumors reached approximately 150 mm^3^ (after another ∼7 days), mice were injected intravenously with 2.5x10^6^ lysing S. typhimurium carrying WT-SVA-NanoLuc RNA. (ii) Weights of mice receiving: CAPPSID-SVA (black) in vaccinated mice, lysing control S. typhimurium in unvaccinated mice (red), or vaccinated mice rechallenged with an additional 2.5x10^6^ SVA viral particles (blue) normalized to initial weight over the first 10 days post-inoculation. (iii) Biodistribution results for (L)(ii) are represented as CFUs/gram of tissue from the spleens, livers, and tumors of mice 3 and 7 days after inoculation with the CAPPSID.

**Figure S3.**
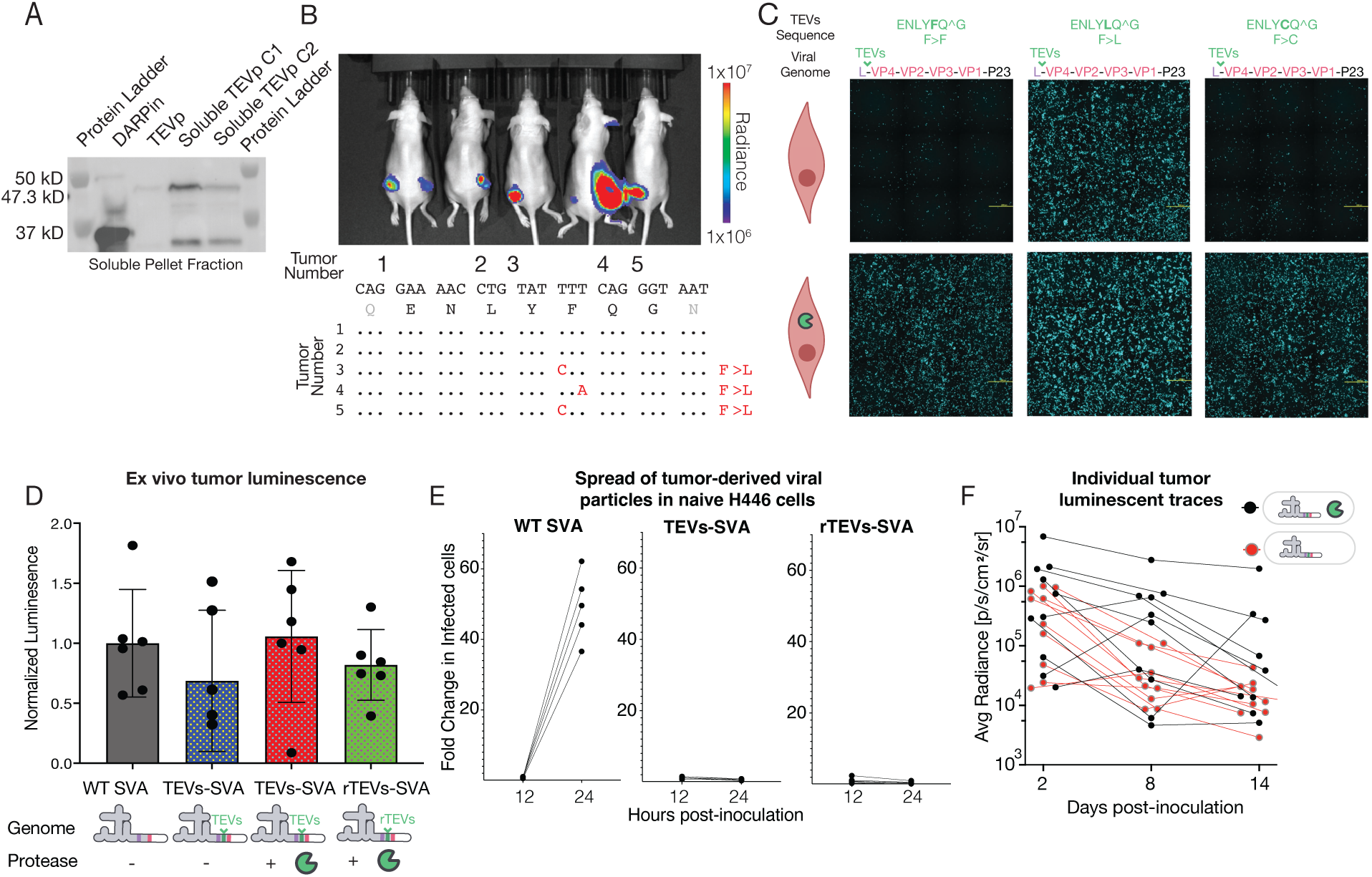
Related to. Figure 4**, Engineered control over viral protein maturation via CAPPSID-delivered protease. (A)** Western blot showing soluble cellular fraction. Columns from left to right: Protein ladder, DARPin expressing Flag-tag (primary ab control), non-optimized TEVp, optimized TEVp clones 1 and 2, and protein ladder for solubility. 43kD band indicates TEV-protease fused to carrier protein. Input was normalized by OD600 values. **(B)** (Top) IVIS of TEVp-dependent delivered by lysing Salmonella with TEVp at day 8 post IT injection. (Bottom) Sequencing results from tumor homogenate of TEVs region viral RNA in the corresponding mice. Dots indicate matching expected bases. **(C)** Transfection of TEVs RNA mutants into cells with (bottom) and without (top) TEVp expression. From left to right, columns are standard TEV site ENLY**F**Q^G; ENLY**L**Q^G site (middle); and ENLY**C**Q^G (right). **(D)** Tumors treated with lysing S. typhimurium carrying WT-SVA-NanoLuc without TEVp, TEVs-SVA-NanoLuc with and without TEVp, and rTEVs-SVA-NanoLuc with TEVp were excised 18 (HPI). Ex-vivo luminescent intensities were quantified with each point representing an individual tumor luminescence divided by the luminescent intensity of the WT-SVA-NanoLuc group. ANOVA evaluation determined no significant difference in the means of groups evaluated. **(E)** Tumor homogenate was inoculated onto naive H446 cells to assay capacity for viral spreading in vitro at 24 hours relative to 12 h measurement. Homogenates were taken from tumors treated IT with bacteria delivering WT-SVA-NanoLuc (left), TEVs-SVA-NanoLuc + TEVp (middle), and rTEVs-SVA-NanoLuc + TEVp (right). 18 hours after IT administration, tumors were isolated, homogenized, freeze-thawed, and clarified to establish resulting viral stocks. 200ul of stock was inoculated on naive H446 cells and imaged at 12h and 24h post-inoculation. Five mice were included per condition. **(F)** Single mouse trajectories of tumor radiance over time. Black lines are mice receiving bacterially-delivered TEVp-dependent virus along with TEVp. Red lines are mice receiving bacterially-delivered TEVp-dependent virus without TEVp.

## References

1 Chen, Y. E. et al. Engineered skin bacteria induce antitumor T cell responses against melanoma. *Science (New York*, N.Y.) 380, 203–210 (2023). 10.1126/science.abp9563

2 Mazzolini, R. et al. Engineered live bacteria suppress Pseudomonas aeruginosa infection in mouse lung and dissolve endotracheal-tube biofilms. Nature Biotechnology 41, 1089–1098 (2023). 10.1038/s41587-022-01584-9

3 Chen, Y. et al. Development of a Listeria monocytogenes-based vaccine against hepatocellular carcinoma. Oncogene 31, 2140–2152 (2012). 10.1038/onc.2011.395

4 Raman, V. et al. Intracellular delivery of protein drugs with an autonomously lysing bacterial system reduces tumor growth and metastases. Nature Communications 12, 6116 (2021). 10.1038/s41467-021-26367-9

5 Camacho, E., Mesa-Pereira, B., Medina, C., Flores, A. & Santero, E. Engineering Salmonella as intracellular factory for effective killing of tumour cells. Scientific Reports 6, 30591 (2016).

6 Mendell, J. R. et al. Single-Dose Gene-Replacement Therapy for Spinal Muscular Atrophy. New England Journal of Medicine 377, 1713–1722 (2017). 10.1056/NEJMoa1706198

7 Russell, S. et al. Efficacy and safety of voretigene neparvovec (AAV2-hRPE65v2) in patients with RPE65-mediated inherited retinal dystrophy: a randomised, controlled, open-label, phase 3 trial. *Lancet (London*, England) 390, 849–860 (2017). 10.1016/S0140-6736(17)31868-8

8 Wickersham, I. R., Finke, S., Conzelmann, K. K. & Callaway, E. M. Retrograde neuronal tracing with a deletion-mutant rabies virus. Nature Methods 4, 47–49 (2007). 10.1038/nmeth999.

9 Desjardins, A. et al. Recurrent Glioblastoma Treated with Recombinant Poliovirus. The New England Journal of Medicine 379, 150–161 (2018). 10.1056/NEJMoa1716435

10 Goetz, C., Dobrikova, E., Shveygert, M., Dobrikov, M. & Gromeier, M. Oncolytic poliovirus against malignant glioma. Future Virology 6, 1045–1058 (2011). 10.2217/fvl.11.76

11 Erickson, A. K. et al. Bacteria Facilitate Enteric Virus Co-infection of Mammalian Cells and Promote Genetic Recombination. Cell host & microbe 23, 77–88.e75 (2017).

12 Robinson, C. M., Jesudhasan, P. R. & Pfeiffer, J. K. Bacterial lipopolysaccharide binding enhances virion stability and promotes environmental fitness of an enteric virus. Cell host & microbe 15, 36–46 (2014).

13 Kuss, S. K. et al. Intestinal microbiota promote enteric virus replication and systemic pathogenesis. *Science (New York*, N.Y.) 334, 249–252 (2011). 10.1126/science.1211057

14 Perez, M. et al. A synthetic consortium of 100 gut commensals modulates the composition and function in a colon model of the microbiome of elderly subjects. Gut Microbes 13, 1–19 (2021). 10.1080/19490976.2021.1919464

15 El Hage, R., Hernandez-Sanabria, E., Calatayud Arroyo, M., Props, R. & Van de Wiele, T. Propionate-Producing Consortium Restores Antibiotic-Induced Dysbiosis in a Dynamic in vitro Model of the Human Intestinal Microbial Ecosystem. Frontiers in Microbiology 10, 1206 (2019). 10.3389/fmicb.2019.01206

16 Li, L. et al. Hydrogel-Encapsulated Engineered Microbial Consortium as a Photoautotrophic “Living Material” for Promoting Skin Wound Healing. ACS applied materials & interfaces 15, 6536–6547 (2023). 10.1021/acsami.2c20399

17 Aalipour, A. et al. Viral Delivery of CAR Targets to Solid Tumors Enables Effective Cell Therapy. Molecular Therapy Oncolytics 17, 232–240 (2020). 10.1016/j.omto.2020.03.018

18 Vincent, R. L. et al. Probiotic-guided CAR-T cells for solid tumor targeting. Science 382, 211–218 (2023). 10.1126/science.add7034

19 Tanoue, T. et al. A defined commensal consortium elicits CD8 T cells and anti-cancer immunity. Nature 565, 600–605 (2019). 10.1038/s41586-019-0878-z

20 Gamboa, L. et al. (bioRxiv, 2021).

21 Khanduja, S. et al. Intracellular delivery of oncolytic viruses with engineered Salmonella causes viral replication and cell death. iScience 27, 109813 (2024). 10.1016/j.isci.2024.109813

22 Roberts, D. M. et al. Hexon-chimaeric adenovirus serotype 5 vectors circumvent pre-existing anti-vector immunity. Nature 441, 239–243 (2006). 10.1038/nature04721

23 Gurbatri, C. R., Arpaia, N. & Danino, T. Engineering bacteria as interactive cancer therapies. *Science (New York*, N.Y.) 378, 858–864 (2022). 10.1126/science.add9667

24 Toso, J. F. et al. Phase I study of the intravenous administration of attenuated Salmonella typhimurium to patients with metastatic melanoma. 20, 142–152 (2002). 10.1200/JCO.2002.20.1.142.

25 Haraga, A., Ohlson, M. B. & Miller, S. I. Salmonellae interplay with host cells. Nature Reviews. Microbiology 6, 53–66 (2008). 10.1038/nrmicro1788

26 Fink, S. L. & Cookson, B. T. Pyroptosis and host cell death responses during Salmonella infection. Cellular Microbiology 9, 2562–2570 (2007). 10.1111/j.1462-5822.2007.01036.x

27 Schoen, C. et al. Bacterial delivery of functional messenger RNA to mammalian cells. Cellular Microbiology 7, 709–724 (2005). 10.1111/j.1462-5822.2005.00507.x

28 Hegazy, W., Xu, X., Metelitsa, L. & Hensel, M. Evaluation of Salmonella enterica type III secretion system effector proteins as carriers for heterologous vaccine antigens. Infection and immunity 80, 1193–1202 (2012).

29 Xu, X., Husseiny, M. I., Goldwich, A. & Hensel, M. Efficacy of intracellular activated promoters for generation of Salmonella-based vaccines. Infection and immunity 78, 4828–4838 (2010).

30 Xu, X., Husseiny, M. I., Goldwich, A. & Hensel, M. Efficacy of intracellular activated promoters for generation of Salmonella-based vaccines. Infection and Immunity 78, 4828–4838 (2010). 10.1128/IAI.00298-10

31 Xiong, G. et al. Novel cancer vaccine based on genes of Salmonella pathogenicity island 2. International Journal of Cancer 126, 2622–2634 (2010). 10.1002/ijc.24957

32 Raman, V. et al. Intracellular delivery of protein drugs with an autonomously lysing bacterial system reduces tumor growth and metastases. Nature Communications 12, 6116 (2021).

33 Xu, X. & Hensel, M. Systematic analysis of the SsrAB virulon of Salmonella enterica. Infection and immunity 78, 49–58 (2009).

34 Xu, X. et al. Development of an Effective Cancer Vaccine Using Attenuated Salmonella and Type III Secretion System to Deliver Recombinant Tumor-Associated Antigens. Cancer research 74, 6260–6270 (2014). 10.1158/0008-5472.CAN-14-1169

35 Juárez-Rodríguez, M. D. et al. Live attenuated Salmonella vaccines displaying regulated delayed lysis and delayed antigen synthesis to confer protection against Mycobacterium tuberculosis. Infection and Immunity 80, 815–831 (2012). 10.1128/IAI.05526-11

36 Manuel, E. R. et al. Enhancement of cancer vaccine therapy by systemic delivery of a tumor-targeting Salmonella-based STAT3 shRNA suppresses the growth of established melanoma tumors. Cancer Research 71, 4183–4191 (2011). 10.1158/0008-5472.CAN-10-4676

37 Husseiny, M. I., Wartha, F. & Hensel, M. Recombinant vaccines based on translocated effector proteins of Salmonella Pathogenicity Island 2. Vaccine 25, 185–193 (2007).

38 Chabloz, A. et al. Salmonella-based platform for efficient delivery of functional binding proteins to the cytosol. Communications Biology 3, 342 (2020).

39 Widmaier, D. M. et al. Engineering the Salmonella type III secretion system to export spider silk monomers. Molecular systems biology 5, 309 (2009).

40 Doamekpor, S. K., Sharma, S., Kiledjian, M. & Tong, L. Recent insights into noncanonical 5’ capping and decapping of RNA. The Journal of Biological Chemistry 298, 102171 (2022). 10.1016/j.jbc.2022.102171

41 Francisco-Velilla, R., Embarc-Buh, A., Abellan, S. & Martinez-Salas, E. Picornavirus translation strategies. FEBS Open Bio 12, 1125–1141 (2022). 10.1002/2211-5463.13400

42 Ansardi, D. C., Porter, D. C., Anderson, M. J. & Morrow, C. D. Advances in Virus Research. Elsevier 46, 1–68 (1996).

43 Din, M. O. et al. Synchronized cycles of bacterial lysis for in vivo delivery. Nature 536, 81–85 (2016).

44 Loessner, H. et al. Remote control of tumour-targeted Salmonella enterica serovar Typhimurium by the use of L-arabinose as inducer of bacterial gene expression in vivo. Cellular Microbiology 9, 1529–1537 (2007). 10.1111/j.1462-5822.2007.00890.x

45 Orta, A. K. et al. The mechanism of the phage-encoded protein antibiotic from ΦX174. *Science (New York*, N.Y.) 381, eadg9091 (2023). 10.1126/science.adg9091

46 Wallace, A. J. et al. E. coli hemolysin E (HlyE, ClyA, SheA): X-ray crystal structure of the toxin and observation of membrane pores by electron microscopy. Cell 100, 265–276 (2000). 10.1016/s0092-8674(00)81564-0

47 Beuzón, C. R. et al. Salmonella maintains the integrity of its intracellular vacuole through the action of SifA. The EMBO journal 19, 3235–3249 (2000). 10.1093/emboj/19.13.3235

48 Reddy, P. S. et al. Seneca Valley virus, a systemically deliverable oncolytic picornavirus, and the treatment of neuroendocrine cancers. J Natl Cancer Inst 99, 1623–1633 (2007). 10.1093/jnci/djm198

49 Burke, M. Oncolytic Seneca Valley Virus: past perspectives and future directions. Oncolytic Virotherapy **Volume** 5, 81–89 (2016).

50 Rudin, C. M. et al. Phase I clinical study of Seneca Valley Virus (SVV-001), a replication-competent picornavirus, in advanced solid tumors with neuroendocrine features. Clin Cancer Res 17, 888–895 (2011). 10.1158/1078-0432.CCR-10-1706

51 Morton, C. L. et al. Initial testing of the replication competent Seneca Valley virus (NTX-010) by the pediatric preclinical testing program. Pediatric Blood & Cancer 55, 295–303 (2010). 10.1002/pbc.22535

52 Poirier, J. T. et al. Characterization of a full-length infectious cDNA clone and a GFP reporter derivative of the oncolytic picornavirus SVV-001. Journal of General Virology 93, 2606–2613 (2012).

53 Geisler, A., Hazini, A., Heimann, L., Kurreck, J. & Fechner, H. Coxsackievirus B3—Its Potential as an Oncolytic Virus. Viruses 13, 718 (2021).

54 Erasmus, J. H. et al. An alphavirus-derived replicon RNA vaccine induces SARS-CoV-2 neutralizing antibody and T cell responses in mice and nonhuman primates. *Science translational medicine*, eabc9396 (2020).

55 Miest, T. S. & Cattaneo, R. New viruses for cancer therapy: meeting clinical needs. Nature Reviews. Microbiology 12, 23–34 (2014). 10.1038/nrmicro3140

56 Palmenberg, A. C. Proteolytic processing of picornaviral polyprotein. Annual Review of Microbiology 44, 603–623 (1990). 10.1146/annurev.mi.44.100190.003131

57 Kapust, R. B., Tözsér, J., Copeland, T. D. & Waugh, D. S. The P1′ specificity of tobacco etch virus protease. Biochemical and Biophysical Research Communications 294, 949–955 (2002).

58 Cabrita, L. D. et al. Enhancing the stability and solubility of TEV protease using in silico design. Protein Science 16, 2360–2367 (2007). 10.1110/ps.072822507

59 van den Berg, S., Löfdahl, P.-A., Härd, T. & Berglund, H. Improved solubility of TEV protease by directed evolution. J Biotechnol 121, 291–298 (2006). 10.1016/j.jbiotec.2005.08.006

60 Aguilar Rangel, M., et al. High-resolution mapping reveals the mechanism and contribution of genome insertions and deletions to RNA virus evolution. Proceedings of the National Academy of Sciences of the United States of America 120, e2304667120 (2023). 10.1073/pnas.2304667120

61 Dougherty, W. G., Cary, S. M. & Parks, T. D. Molecular genetic analysis of a plant virus polyprotein cleavage site: a model. Virology 171, 356–364 (1989). 10.1016/0042-6822(89)90603-x

62 Zhang, Z. et al. Development of an Agrobacterium-delivered CRISPR/Cas9 system for wheat genome editing. Plant Biotechnol J 17, 1623–1635 (2019). 10.1111/pbi.13088

63 Schaffner, W. Direct transfer of cloned genes from bacteria to mammalian cells. Proceedings of the National Academy of Sciences 77, 2163–2167 (1980).

64 Narayanan, K. & Warburton, P. E. DNA modification and functional delivery into human cells using Escherichia coli DH10B. Nucleic Acids Research 31, e51 (2003). 10.1093/nar/gng051

65 Grillot-Courvalin, C., Goussard, S., Huetz, F., Ojcius, D. M. & Courvalin, P. Functional gene transfer from intracellular bacteria to mammalian cells. Nature Biotechnology 16, 862–866 (1998).

66 Grillot-Courvalin, C., Goussard, S. & Courvalin, P. Wild-type intracellular bacteria deliver DNA into mammalian cells. Cellular Microbiology 4, 177–186 (2002). 10.1046/j.1462-5822.2002.00184.x

67 Kong, W. et al. Regulated programmed lysis of recombinant Salmonella in host tissues to release protective antigens and confer biological containment. Proceedings of the National Academy of Sciences 105, 9361–9366 (2008).

68 Kong, W., Brovold, M., Koeneman, B. A., Clark-Curtiss, J. & Curtiss, R. Turning self-destructing Salmonella into a universal DNA vaccine delivery platform. Proceedings of the National Academy of Sciences of the United States of America 109, 19414–19419 (2012). 10.1073/pnas.1217554109

69 Xiang, S., Fruehauf, J. & Li, C. J. Short hairpin RNA-expressing bacteria elicit RNA interference in mammals. Nature Biotechnology 24, 697–702 (2006). 10.1038/nbt1211

70 Guo, H. et al. Targeting tumor gene by shRNA-expressing Salmonella-mediated RNAi. Gene Therapy 18, 95–105 (2011). 10.1038/gt.2010.112

71 Weiss, S. & Chakraborty, T. Transfer of eukaryotic expression plasmids to mammalian host cells by bacterial carriers. Current Opinion in Biotechnology 12, 467–472 (2001). 10.1016/s0958-1669(00)00247-0

72 Acevedo, A., Brodsky, L. & Andino, R. Mutational and fitness landscapes of an RNA virus revealed through population sequencing. Nature 505, 686 (2014).

73 Drake, J. W. Rates of spontaneous mutation among RNA viruses. Proceedings of the National Academy of Sciences of the United States of America 90, 4171–4175 (1993).

74 Sanjuán, R., Nebot, M. R., Chirico, N., Mansky, L. M. & Belshaw, R. Viral mutation rates. Journal of Virology 84, 9733–9748 (2010). 10.1128/JVI.00694-10

75 Domingo, E., García-Crespo, C., Lobo-Vega, R. & Perales, C. Mutation Rates, Mutation Frequencies, and Proofreading-Repair Activities in RNA Virus Genetics. Viruses 13, 1882 (2021). 10.3390/v13091882

76 Singer, Z. S., Ambrose, P. M., Danino, T. & Rice, C. M. Quantitative measurements of early alphaviral replication dynamics in single cells reveals the basis for superinfection exclusion. Cell Systems 12, 210–219.e213 (2021). 10.1016/j.cels.2020.12.005

